# Enhancing NosZ Activity to Reduce N_2_O Emissions from Biological Wastewater Treatment Systems

**DOI:** 10.1101/2024.09.22.614384

**Authors:** Xueyang Zhou, Bharat Manna, Boyu Lyu, Naresh Singhal

## Abstract

Nitrous oxide (N_2_O) emissions from wastewater treatment plants, with a warming potential 298 times that of CO_2_, pose a significant challenge to lowering their carbon footprint. Current mitigation strategies focus on limiting N_2_O formation during nitrification and denitrification but overlook microbial reduction mechanisms. This study examines the potential for enhancing nitrous oxide reductase (NosZ) activity to reduce N_2_O to N_2_. We hypothesize that strategic oxygen manipulation can enhance N_2_O destruction by continuous NosZ expression and enable NosZ activation in microorganisms with superior NosZ capabilities. We assess microbial community function and metabolic regulation using metagenomics and metaproteomics to clarify the effect of intermittent aeration regimes on N_2_O emission. Intermittent aeration with periodic anoxic exposure significantly reduced N_2_O emissions with 71% nitrogen removal by enhancing the metabolic activity of *Hyphomicrobium*. NosZ activity increased by 4- to 6.5-fold after system adaptation to oxygen modulations, compared to continuous oxic-anoxic cycling without the anoxic phase. The latter resulted in increased N_2_O emissions due to suppressed NosZ activity and higher N_2_O production from *Methylobacillus*, which uses nitric oxide as an alternative electron acceptor. Our finding that strategic oxygen manipulation can energize N_2_O destruction lays the foundation for developing next-generation wastewater treatment technologies for mitigating N_2_O emissions.

## INTRODUCTION

Nitrous oxide (N_2_O) emissions from biological nitrogen removal processes, which have a global warming potential 298 times greater than carbon dioxide, pose a significant challenge for wastewater treatment plants aiming to achieve carbon neutrality^1–3^. Despite extensive research, current N_2_O mitigation strategies remain constrained by a production-centric focus, overlooking the transformative potential of microbial reduction mechanisms^4–6^. In contrast to managing N_2_O production during nitrification and denitrification^7–10^, the true potential for mitigation lies in the biological destruction pathway—specifically, the reduction of N_2_O to N_2_ by nitrous oxide reductase (NosZ), the only enzyme capable of complete N_2_O elimination^11–15^. This research challenges conventional strategies^1,4,16–19^ by exploring how targeted deoxygenation can unlock the latent N_2_O destruction capabilities of microbial communities^20^.

Recent studies reveal a nuanced interplay between oxygen availability and N_2_O reduction, highlighting the metabolic flexibility of denitrifying microorganisms^21,22^. Historical oxygen manipulation studies show mixed results^23^, with some reducing N_2_O emissions^4,5,24,25^ and others increasing them^26,27^. These inconsistencies underscore a critical knowledge gap in understanding the mechanisms linking oxygen variation to N_2_O dynamics^4,5,28–30^. We hypothesize that strategic oxygen manipulation can enhance N_2_O destruction through three key mechanisms: (i) maintaining continuous NosZ expression, (ii) enabling NosZ activation, and (iii) selectively enhancing microorganisms with superior NosZ capabilities. This approach shifts from traditional emission management to active microbial-driven N_2_O elimination.

Our study investigates microbial community function and metabolic regulation under controlled intermittent aeration regimes. We aim to identify key microbial contributors, their enzymatic pathways, and the regulatory mechanisms governing N_2_O production and reduction under varying oxygen conditions. By bridging fundamental microbial mechanisms with practical applications, we aim to provide an evidence-based framework for developing next-generation wastewater treatment technologies for effectively mitigating N_2_O emissions from biological wastewater treatment processes.

## MATERIALS AND METHODS

### Bioreactor Operation

Activated sludge used in this study was sourced from the Māngere Wastewater Treatment Plant in Auckland, New Zealand. Three identical cylindrical acrylic bioreactors, each with a 1 L working volume, were operated simultaneously under varying aeration conditions at 20⏢. The same activated sludge was used in triplicate to ensure biological consistency. Each 1 L reactor contained 2.75 g/L of mixed liquor-suspended solids and was supplemented with 3.84 g/L of NaHCO3 as an inorganic carbon source, along with 1 mL of a trace element solution (Table S1). Over a 48-hour period, 50 mL of concentrated artificial wastewater (Table S2) was continuously fed into each reactor at an equal rate using a syringe pump, while magnetic stirring ensured uniform mixing. The bioreactors were operated under three aeration modes, aimed at suppressing denitrification, inducing N_2_O production, or promoting N_2_O reduction, respectively. Each mode was tested at two oxygen levels, resulting in six conditions: constant aerobic (CA) with steady oxygen levels of 2 and 8 mg/L (Figure 1a, 1b); continuous perturbed (CP) with cyclic oxygen variations of 0-2 and 0-8 mg/L without intermittent anoxic stage (Figure 1c, 1d); and, intermittent perturbed (IP) with cyclic oxygen variations that include an anoxic stage (Figure 1e, 1f).

**Figure 1.**
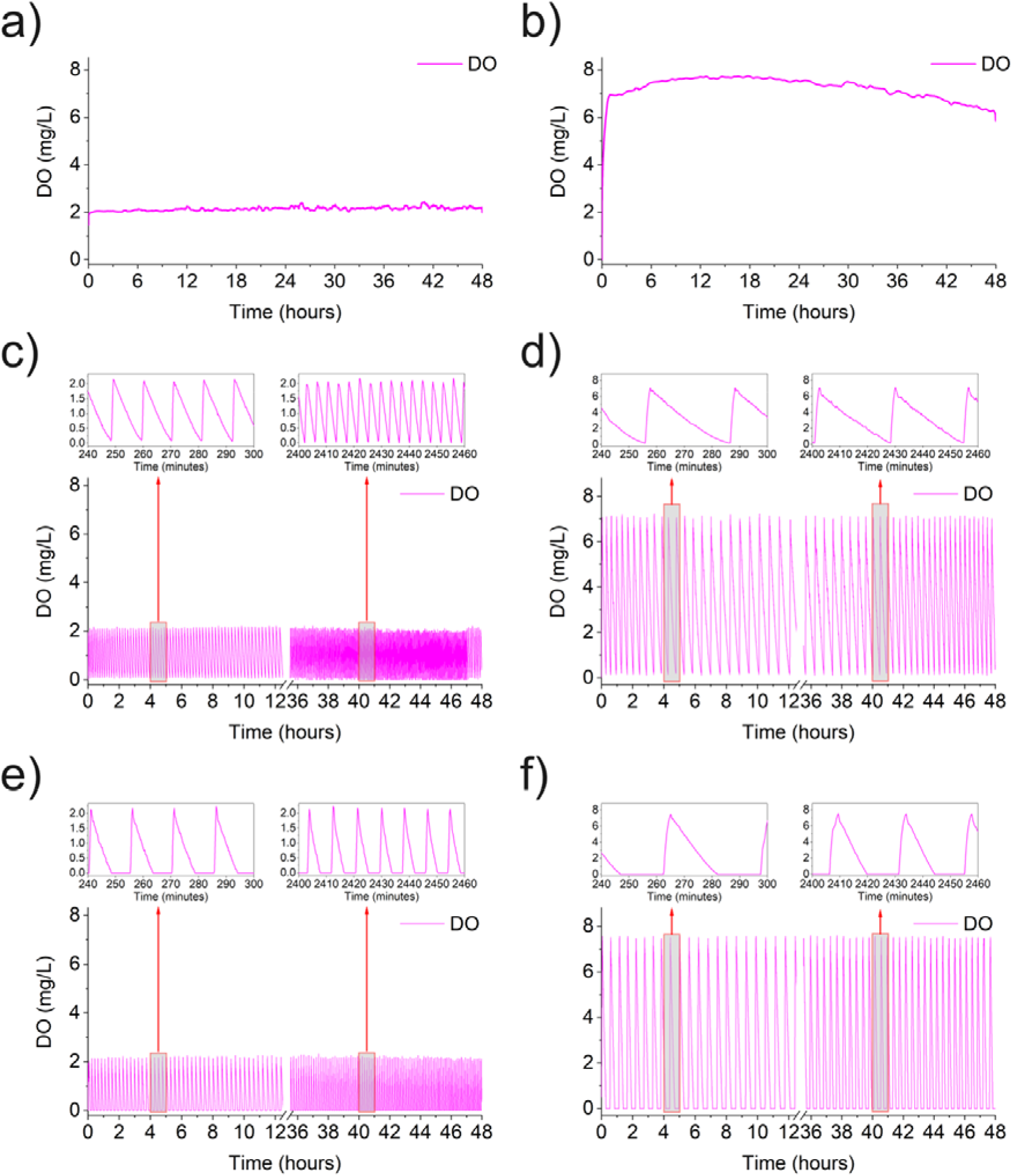
Dissolved oxygen (DO) concentration profiles under six aeration conditions: CA2, CA8, CP2, CP8, IP2, and IP8. Each plot highlights distinct modes of oxygen availability, showcasing constant aerobic (CA), continuous perturbed (CP), and intermittent perturbed (IP) modes at low (2 mg/L) and high (8 mg/L) DO levels.

### N_2_O Emissions

Gaseous N_2_O emissions were quantified using gas chromatography (Shimadzu GC-BID/FID 2010 Plus), and dissolved N_2_O concentrations were monitored with online N_2_O sensors (Unisense, Denmark). Detailed analytical methods, including sensor calibration, gas sampling, and data processing procedures, are provided in Supplementary Materials S1.3.

### In-situ N_2_O reductase (N_2_OR) Activity

In-situ N_2_OR activity was assessed by introducing a saturated N_2_O solution into the bioreactor and monitoring the consumption of dissolved N_2_O using an online sensor (Unisense, Denmark). The N_2_O consumption rate was determined by analyzing the time-dependent decrease in dissolved N_2_O concentrations.

### Metagenomics

DNA was extracted from sludge samples using the DNeasy PowerSoil Kit (Qiagen, Germany) following the manufacturer’s protocol. Samples were collected at the start of the experiment (six biological replicates) for nitrogen transformer identification and at the 48-hour mark under dissolved oxygen (DO) 2 and DO8 conditions to construct the metaproteomics library. The extracted DNA underwent Illumina sequencing on a HiSeq platform. Details on sample preparation, sequencing procedures, and bioinformatics analysis using SqueezeMeta^31^ are provided in Supplementary Materials S1.4.

### Metaproteomics

Protein extraction and analysis were conducted for sludge samples collected under different aeration conditions. Biological replicates included five from the start of the experiment, three from 48-hour DO2, and two from 48-hour DO8 conditions. Proteins were extracted, purified, digested, and analyzed via nano LC-MS/MS using a TripleTOF 6600 mass spectrometer. Protein identification was conducted against a database established by metagenomics results of the samples using MetaProteomeAnalyzer version 3.4^32^. Full details of the experimental protocol are available in Supplementary Materials S1.5.

## RESULTS

### Nitrogen Removal Under Perturbed Aeration

Under CA conditions at both tested DO concentrations (CA2 and CA8), nitrogen removal was minimal (<3%), indicating the predominance of aerobic metabolic pathways (Figure S1, S2). CP conditions (CP2 and CP8) facilitated limited denitrification, resulting in approximately 8% conversion of nitrogen to gaseous forms (Figure S3, S4). In contrast, IP conditions (IP2 and IP8) enabled significant nitrogen removal (∼71%), driven by enhanced denitrification through strategic oxygen availability patterns (Figure S5, S6).

### Role of Oxygen Modulation in N_2_O Dynamics

To understand the role of oxygen modulation in N_2_O dynamics, we analyzed dissolved and gaseous N_2_O emissions under three aeration modes—CA, CP, and IP. The N_2_O emissions show distinct trends in dynamics (Figure 2). IP showed lower dissolved N_2_O accumulation and stabilized gaseous emissions than CP, which was comparable to the N_2_O emissions under CA. N_2_O transformation dynamics exhibited three characteristic phases (Figure 2): an initial accumulation phase (Phase I), a depletion phase (Phase II), and lastly a stabilization phase (Phase III). During Phase I, the dissolved N_2_O concentrations increased across all aeration modes and showed significant variation among the observed rates for different conditions at 95% confidence (Table S3). The dissolved N_2_O accumulation rates during the initial phase were significantly lower under IP (0.034 mg/L/h for IP2 and 0.007 mg/L/h for IP8) compared to CP (0.064 mg/L/h for CP2 and 0.089 mg/L/h for CP8). Gaseous N_2_O emissions mirrored these trends, with substantially lower N_2_O-N emitted in the first 8 hours under IP (5.99 mg N and 5.08 mg N under IP2 and IP8, respectively) than CP (9.32 mg N and 12.07 mg N under CP2 and CP8, respectively). This reduction in initial accumulation suggests that anoxic intervals inherent to IP strategies effectively limit N_2_O production pathways. During Phase II, the N_2_O emissions decreased, in particular for CP2, CP8, and IP2. The depletion rates under IP2 (-0.022 mg/L/h) were comparable to CP (-0.023 mg/L/h for CP2 and -0.039 mg/L/h for CP8); however, IP consistently showed lower gaseous emissions throughout this phase (0.73 to 0.21 mg N/hour under IP2 versus 1.94 to 0.29 mg N/hour under CP2). This trend continued until stabilization was achieved. Phase III data show that while the dissolved N_2_O concentrations were lowest under IP (0.11–0.26 mg/L for IP2 and 0.036–0.12 mg/L for IP8), gaseous emissions became negligible across all conditions, signaling an equilibrium between N_2_O production and reduction processes. Moreover, NosZ expression and activation, leading to higher N_2_OR activity and dissolved N_2_O consumption rates, were highest under IP (1.23 mg N_2_O-N/L/min and 4.19 mg N_2_O-N/L/min for IP2 and IP8, respectively) and showed 4- to 6.5-fold exceedance over CP conditions with the same oxygen range (Figure 3, Table S4).

**Figure 2.**
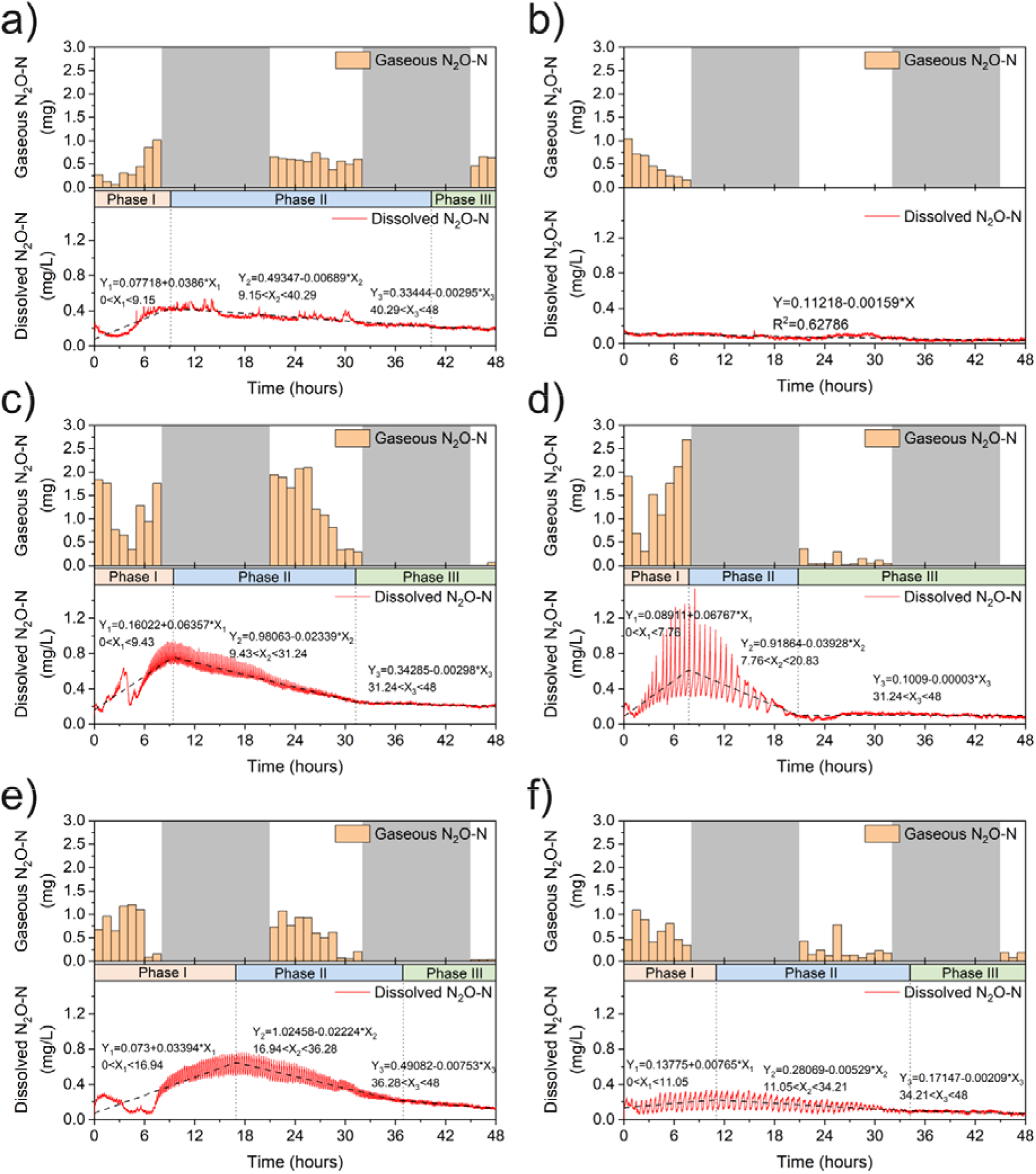
Gaseous and dissolved N_2_O concentration dynamics under six aeration conditions: CA2, CA8, CP2, CP8, IP2, and IP8. Gaseous N_2_O emissions were measured during 0–8 hours, 22–32 hours, and 46–48 hours, while dissolved N_2_O concentrations were continuously monitored over 48 hours.

**Figure 3.**
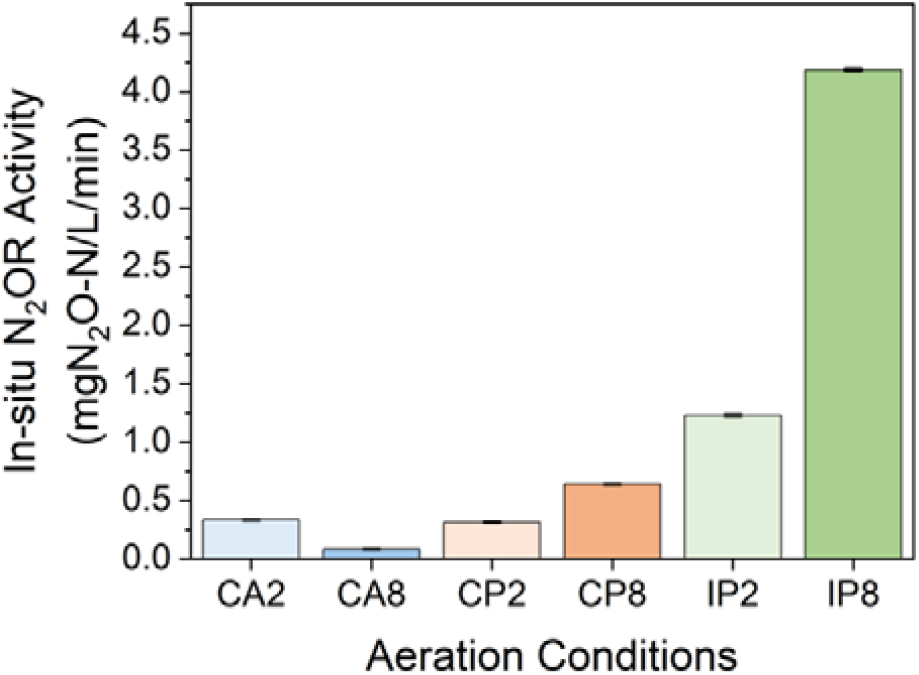
In situ N_2_O reductase (N_2_OR) activity, measured as the rate of dissolved N_2_O consumption after 48 hours of exposure to six aeration conditions. Results indicate enhanced N_2_OR activity under IP conditions compared to CA and CP modes.

### Microbial Contributors to N_2_O Emissions

Metagenomic and metaproteomic analyses (Figure 4, Figure S7) identified the key organisms responsible for N_2_O dynamics and how aeration strategies influence their metabolic activities. *Nitrosomonas* is the dominant ammonia-oxidizing genus that catalyzes the conversion of ammonia to hydroxylamine and subsequently to nitrite, and *Nitrospira* is the primary nitrite-oxidizing genus for nitrite to nitrate oxidation (Figure 4); both possess genes for partial denitrification pathways to convert nitrite to nitric oxide (NO) and N_2_O, potentially contributing to N_2_O emissions. *Methylobacterium* and *Hyphomicrobium* stand out as the two main denitrifying genera (Figure 4) but possess different denitrification functional genes. *Methylobacterium* contains genes for NO and N_2_O production and lacks genes for N_2_O reduction, making it a potential source of N_2_O production. In contrast, *Hyphomicrobium* has a complete set of denitrification-related genes, which includes the *nosZ* gene, and could contribute to N_2_O reduction.

**Figure 4.**
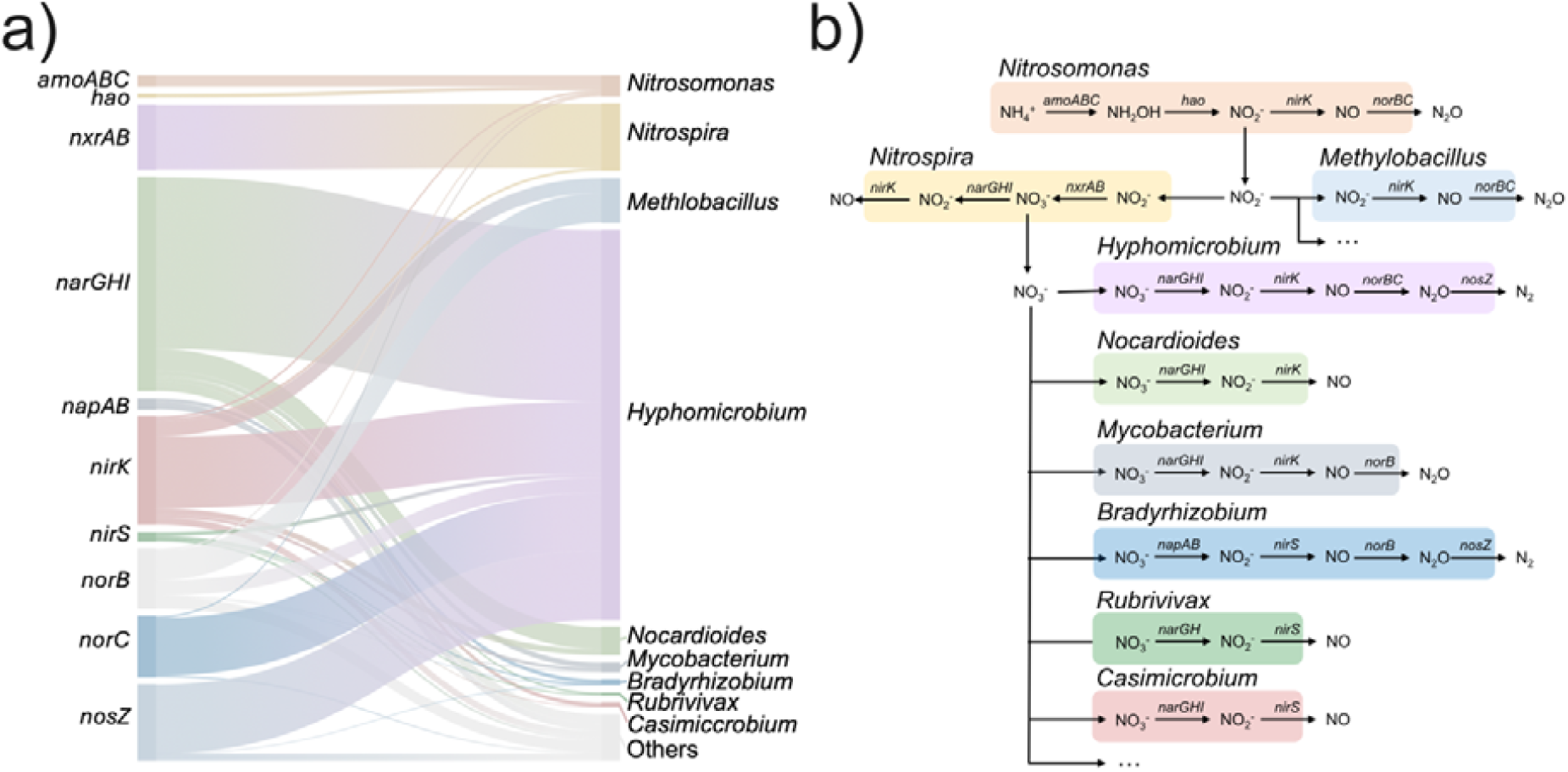
Metagenomic insights into microbial community structure and functional differentiation of nitrogen-transforming genera. (a) Source microorganisms and their contribution to nitrogen transformation-related genes. (b) Functional gene differentiation among nitrogen-transforming genera.

An enzymatic analysis showed significant differences in the abundances of denitrification enzymes under different aeration conditions (Figure S7). Nitric oxide reductase (NorB) and NosZ emerged as critical enzymes driving the N_2_O dynamics. NorB, responsible for N_2_O production, showed higher abundance under CP, compared to CA and IP, and its activity was primarily attributed to *Methylobacterium*, a genus within the class Betaproteobacteria (Figure 5). Conversely, NosZ, which facilitates N_2_O reduction, was significantly higher under IP and is primarily associated with *Hyphomicrobium* within the class Alphaproteobacteria (Figure 5). The differences in enzyme abundance under different aeration conditions mirror the observed N_2_O emissions. Since N_2_O producers and reducers tend to favor different aeration modes, oxygen availability under some aeration patterns could influence the interactions between N_2_O producers and reducers, resulting in dynamic changes in the dissolved N_2_O pool.

**Figure 5.**
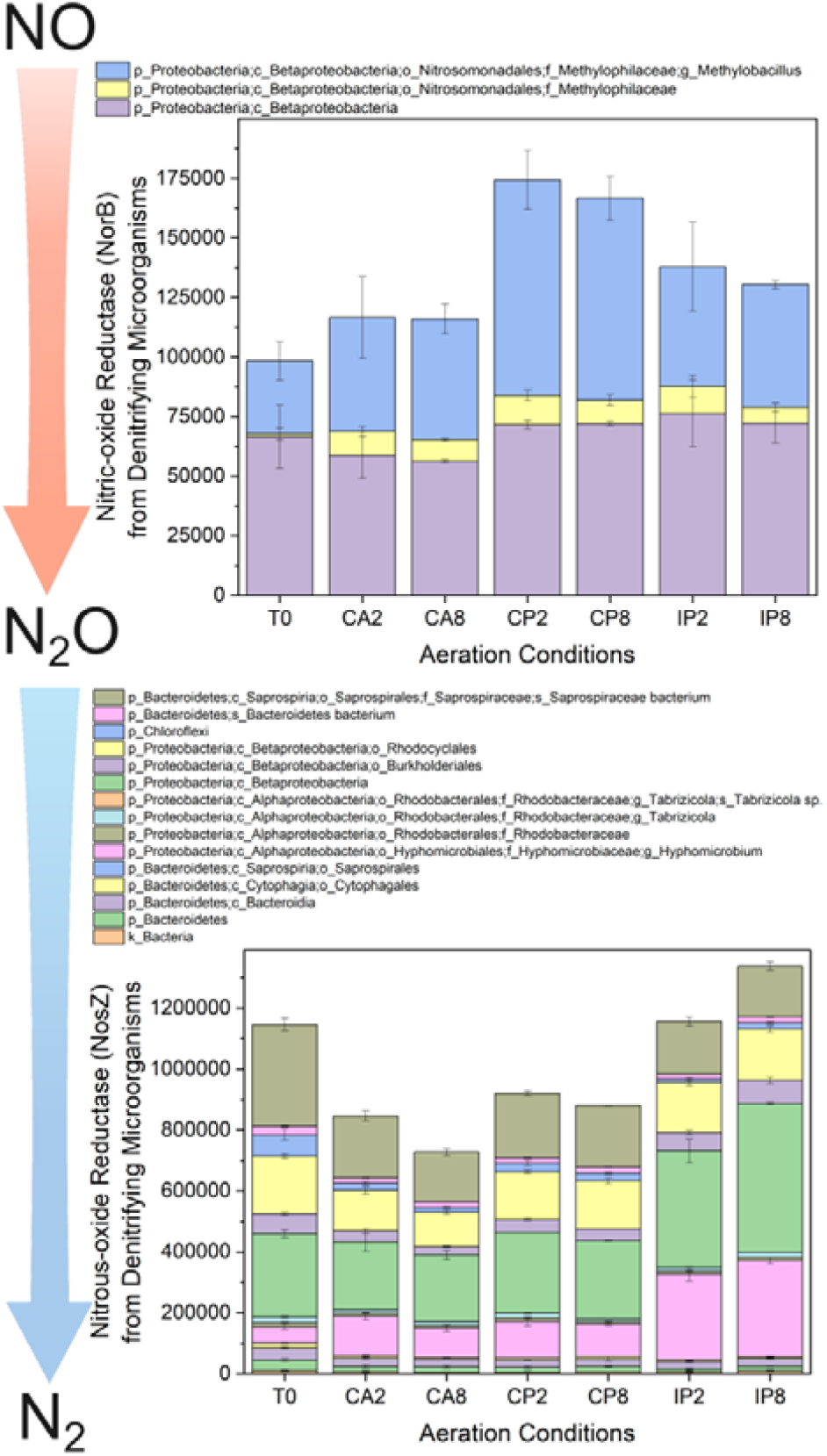
Microbial sources of nitric oxide reductase (NorB) and nitrous oxide reductase (NosZ) identified through metaproteomic analysis. Contributions under CP and IP conditions illustrate distinct microbial drivers for N_2_O production and reduction.

### Mechanisms of N_2_O Metabolism

A detailed metaproteome analysis revealed distinct adaptation strategies employed by *Methylobacillus* and *Hyphomicrobium* under different aeration patterns. Increasing the oxygen level from 2 to 8 mg/L did not significantly affect these strategies.

#### Methylobacillus Adaptations

Under CP conditions, *Methylobacillus* exhibited higher expression of cytochrome c oxidase cbb3-type and NorB, indicating that N_2_O formation occurs as a metabolic byproduct under limited oxygen availability (Figure 6a). This adaptation was supported by stabilized NO concentrations of approximately 14 μM (Figure S8), enabling sustained electron transport chain function through NO utilization. The absence of NosZ in *Methylobacillus* resulted in its inability to reduce the N_2_O it produced. Two-way ANOVA analysis (Figure S9) showed that aeration mode significantly influenced *Methylobacterium* metabolism (p = 0.003), while oxygen level had an insignificant effect (p = 0.74). Overall enzyme expression in *Methylobacterium* declined under IP conditions compared to CA and CP conditions, contributing to the suppression of N_2_O production (Figure S10).

**Figure 6.**
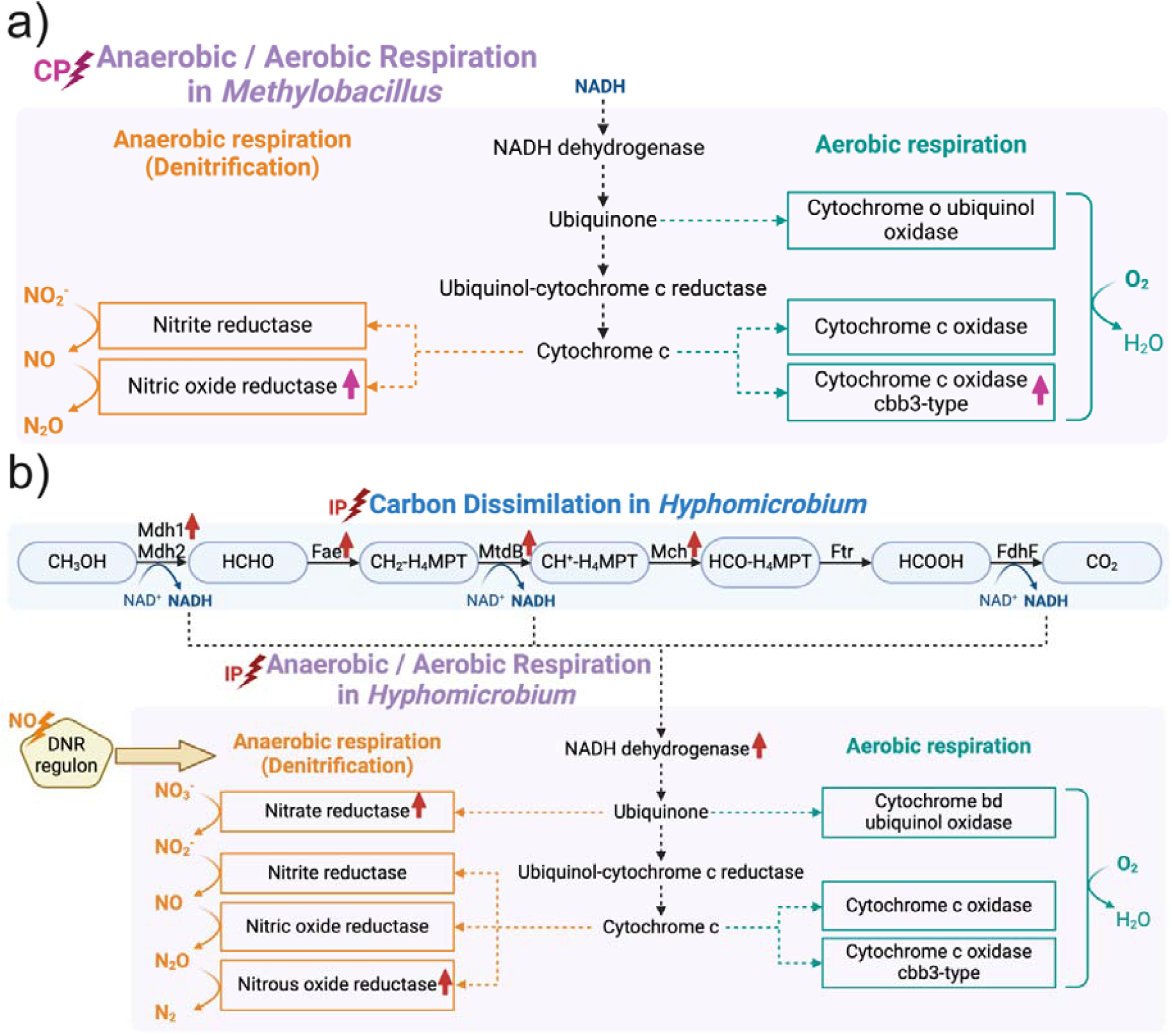
Metabolic regulatory mechanisms of N_2_O-related processes in (a) *Methylobacillus* under CP conditions, characterized by upregulation of nitric oxide reductase and cytochrome c oxidase cbb3-type, and (b) *Hyphomicrobium* under IP conditions, showing enhanced electron generation and upregulation of nitrate reductase and nitrous oxide reductase.

#### Hyphomicrobium Adaptations

*Hyphomicrobium* exhibited elevated enzyme expression ability under IP conditions (p = 0.008) (Figure S11), contrasting with the metabolic response patterns observed in *Methylobacterium*. This indicates that *Hyphomicrobium* possesses a distinct adaptive mechanism to redox fluctuations, including a strong potential for reducing N_2_O (Figure S12). Under IP conditions, *Hyphomicrobium* demonstrated comprehensive metabolic reprogramming (Figure 6b). Enhanced expression of carbon dissimilation enzymes included upregulation of methanol dehydrogenase (Mdh), methylene-tetrahydromethanopterin dehydrogenase (MtdB), and NADH dehydrogenase. Upregulation of nitrate reductase (NarGHJI) and NosZ enhanced energy production and N_2_O reduction capabilities under low-oxygen conditions. A key regulatory adaptation in *Hyphomicrobium* involved upregulating the CRP/FNR family transcriptional regulator, specifically the dissimilatory nitrate respiration regulator (DNR) (Figure 6b). This regulator responded to elevated NO concentrations (∼21 μM) (Figure S8), coordinating terminal enzyme expression patterns. Integrating these adaptive mechanisms enabled *Hyphomicrobium* to effectively utilize NO as a signal, optimizing anaerobic respiration and N_2_O reduction under IP conditions.

## DISCUSSION

Our investigation into strategic oxygen manipulation in wastewater treatment systems has revealed complex mechanisms governing N_2_O transformation. Detailed analyses of N_2_O dynamics, enzyme regulation, and metabolic adaptations provide strong evidence supporting our hypothesis that controlled oxygen availability can enhance community-wide N_2_O destruction through three interconnected mechanisms.

### Continuous NosZ Expression

Contrary to conventional assumptions about oxygen sensitivity, we demonstrated that *Hyphomicrobium* maintains continuous NosZ expression under specific intermittent aeration (IP conditions). Stable enzyme levels were exhibited after the system adapted to the aerobic-anoxic transition, suggesting metabolic resilience and adaptability essential for effective N_2_O mitigation. This finding aligns with observations from other systems, including mixed microbial communities (e.g., soil microorganisms^33^ and permeable sediments^34^) and specific pure cultures (e.g., *Azospira sp*.^22^, *Thauera sp*.^35^, and *Pseudomonas stutzeri*^36^), where NosZ expression occurs due to relaxed regulation of respiratory genes by oxygen and metabolic flexibility to exploit periodic supplies of electron acceptors under fluctuating oxygen conditions^37–39^.

### Enhanced NosZ Activation

Limiting oxygen by introducing an anoxic phase in the IP cycles significantly enhanced NosZ activation, resulting in markedly higher N_2_O reduction rates under IP conditions compared to CA and CP conditions. This suggests a sophisticated regulatory mechanism in *Hyphomicrobium*, similar to the resilience seen in *Azospira sp*., which demonstrates rapid recovery of NosZ activity after oxygen exposure^22,40^. Additionally, evidence suggests that denitrification proteins expressed under oxic conditions remain functional, further supporting this adaptability^34^. This tolerance ensures sustained carbon metabolism and continuous electron flow for energy production, regardless of oxygen availability^5,41,42^, highlighting NosZ’s critical role in maintaining robust N_2_O reduction under fluctuating environmental conditions^43,44^.

### Selective Enhancement of NosZ-Containing Microorganisms

The oxygen manipulation strategy selectively enhanced microorganisms with superior NosZ capabilities, particularly *Hyphomicrobium*, which leveraged its complete denitrification pathway and metabolic flexibility to sustain robust N_2_O reduction even under fluctuating oxygen conditions^39,45^. In contrast, *Methylobacillus*, which operates with an incomplete denitrification pathway, relied on cbb3-type cytochrome c oxidase and NorB to maximize the utilization of electron acceptors^46^. However, this adaptation led to increased N_2_O production, highlighting the drawbacks of incomplete denitrifiers in dynamic environments^47,48^.

### Ecological Dynamics and Community-Wide Impact

The contrasting responses of these microorganisms emphasize the importance of targeting complete denitrifiers in optimizing N_2_O mitigation strategies. The shared utilization of denitrification intermediates among microbial communities suggests an ecological advantage in enhancing nosZ-containing species^45^. Given that the potential for N_2_O reduction often surpasses its production capacity^20^, fostering these organisms can amplify their collective impact, creating a synergistic effect that reduces N_2_O emissions and contributes to the overall efficiency of nitrogen cycling within the microbial ecosystem^49^.

### Implications for N_2_O Mitigation Strategies

Implementing strategies solely targeting N_2_O production may be suboptimal^50^. While stable oxygen conditions (CA) effectively suppress N_2_O production, they lack resilience to sudden oxygen fluctuations, often leading to abrupt N_2_O surges^9,51,52^. In contrast, specific IP conditions have been shown to offer superior ways of reducing N_2_O emissions^4,5,24^. Our study provides significant evidence that enhancing the N_2_O reduction capacity of nosZ-containing microorganisms plays a critical role in buffering against N_2_O accumulation, reinforcing the effectiveness of IP strategies. These findings underscore the importance of designing mitigation approaches that address both N_2_O production and reduction mechanisms^53,54^. Targeted enhancement of N_2_O destruction represents a promising pathway to optimize mitigation efforts and maintain stable system performance under variable operational conditions. This research establishes a comprehensive mechanistic framework for controlling N_2_O emissions in biological nitrogen removal systems, demonstrating the effectiveness of strategic oxygen manipulation in promoting NosZ activity for N_2_O reduction.

## Associated content Supporting information

Detailed materials and methods, the performance of biosystems, N_2_O accumulation dynamics, N_2_O consumption rates, the abundance of denitrification-related enzymes, NO concentrations, and enzyme expression analysis (PDF)

## Author information Corresponding authors

*Phone: +64 9 923 4512; fax: +64 9 373 7462.

## Funding sources

The study was supported by the Marsden Fund Council from Government funding, managed by Royal Society Te Apārangi, and a Smart Ideas grant from the Endeavour Fund managed by the Ministry for Business, Innovation & Employment Hīkina Whakatutuki.

## Acknowledgments

We thank Nikki Freed for the assistance with metagenomic sequencing. We thank Martin Middleditch and George Guo for their assistance with metaproteomics analysis. We also thank Watercare Services Limited for providing the activated sludge culture from the Māngere Wastewater Treatment Plant. The authors acknowledge assistance from the New Zealand eScience Infrastructure (NeSI) high-performance computing facilities and the Centre for eResearch at the University of Auckland.

## Notes

### Competing Interest Statement

The authors have declared no competing interest.

### Summary of Updates

The manuscript has been revised to better describe the experimental phenomena and elucidate the mechanism.

