## Supplementary for "Enhancing NosZ Activity to Reduce N_2_O Emissions from Biological Wastewater Treatment Systems"

**for**

**S1. SUPPLEMENTARY METHODS**

**S1.1. Bioreactor operation and performance**

### Table S1. Composition of the trace element solution

| **Chemical** | **Concentration (g/L)** |
| --- | --- |
| EDTA | 2.50 |
| ZnSO_4_·7H_2_O | 1.10 |
| CoCl_2_·6H_2_O | 0.80 |
| MnCl_2_·4H_2_O | 2.55 |
| MgSO_4_·7H_2_O | 20.0 |
| CuSO_4_·5H_2_O | 0.86 |
| (NH_4_)_6_Mo_7_O_24_·4H_2_O | 0.07 |
| CaCl_2_·2H_2_O | 2.75 |
| FeSO_4_·7H_2_O | 2.57 |

### Table S2. Composition of the concentrated artificial wastewater

| **Chemical** | **Concentration** |
| --- | --- |
| Methanol (mL/L) | 26.94 |
| NH_4_Cl (g/L) | 24.46 (IP); 30.57 (CP or CA) |
| KH_2_PO_4_ (g/L) | 2.76 |
| K_2_HPO_4_ (g/L) | 2.76 |
| NaHCO_3_ (g/L) | 76.80 |

**S1.2. Water Quality Analysis**

Chemical oxygen demand (COD) was analysed using COD LR reagent vials (Hach, USA). NH_4_^+^-N concentration was measured with an AmVer Test N Tube Reagent Set (Hach, USA). NO_2_^-^-N and NO_3_^-^-N concentrations were measured using an ICS2100 Ion Chromatograph (ThermoFisher, USA).

**S1.3. N_2_O Emission Measurement**

In this study, the measurement of gaseous N_2_O emissions involved collecting gases released during the reactor operation using multiple gas sampling bags with a maximum volume of 25 L. After collection, the gas samples were analyzed for N_2_O concentrations using a Shimadzu GC-BID/FID 2010 plus gas chromatography. The chromatographic column employed was an HP-Plot/molecular sieve (29.8 m × 0.53 mm inner diameter × 40 μm film), and the detection method included an injection port temperature of 150℃, oven temperature of 160℃, and BID detector temperature of 180℃. The N_2_O-N content (mg) released into the air from the reactor was calculated based on the N_2_O concentration in the gas sample and the volume of gas exiting the reactor.

For the measurement of dissolved N_2_O concentrations (mg/L) in the reactor system, we utilized N_2_O online sensors (Unisense, Denmark), following the manufacturer's guidelines. To ensure accurate and reliable results, the sensors were calibrated and maintained according to the manufacturer's recommendations. Moreover, the data collected from the sensors were processed piecewise linear fitting using Origin Pro 2024.

**S1.4. Metagenomics**

For DNA extraction, 10 mL sludge samples were collected. The extraction process was executed according to the protocol of DNeasy PowerSoil Kit (Qiagen, German), and the extracted DNA was stored at -20°C for subsequent analysis.

The extracted DNA samples were then dispatched to the Auckland Genomics Centre for metagenomic analyses. The laboratory procedures included concentrating the DNA samples, performing a quality control (QC) check, and preparing the DNA for Illumina sequencing, which included PCR and indexing. Following the library QC, normalization was performed, and a final pool check was conducted using a bioanalyzer. Finally, sequencing was performed on a HiSeq platform, generating approximately 400 million 2x150bp paired-end reads.

We performed preprocessing steps to eliminate low-quality sequences using Trimmomatic v0.39^1^ with specific quality thresholds as follows: LEADING:3, TRAILING:3, SLIDINGWINDOW:10:15, and MINLEN:50 with the following settings: TruSeq3-PE-2.fa:2:30:10:2:keepBothReads^2^. For metagenomic taxonomic and functional profiling, we employed the SqueezeMeta v1.5.2 pipeline^3^. Co-assembly was performed using Megahit v1.2.9^4^, and short contigs (<200 bps) were filtered out using prinseq v0.20.4^5^. Within the SqueezeMeta pipeline, the Barrnap v0.9^6^ tool was used for predicting RNAs, while Prodigal v2.6.3^7^ was utilized for predicting ORFs. Taxonomic ranks were assigned against the NCBI GenBank nr database^8^ with identify thresholds of 85, 60, 55, 50, 46, 42, and 40% for the species, genus, family, order, class, phylum, and superkingdom ranks, respectively^9^. Kyoto Encyclopaedia of Genes and Genomes^10^ was used for functional assignments with Diamond v2.0.14^11^ with a maximum e-value threshold of 1 × 10^−3^ and a sequence identify threshold of 50%. Bowtie2 v2.3.4.1^12^ was used for read mapping against the contigs. Both functional genes and ARGs were normalized to transcripts per million (TPM). The metagenomics data were analysed using the SQMtools R package v1.6.3^13^.

**S1.5. Metaproteomics**

The 5 ml sample was collected from the bioreactor at the end of the reaction, then pelleted, washed, and centrifuged at 6,113 x g for 5 minutes at 4 °C. The pellets were rapidly frozen in liquid nitrogen and preserved at -80 °C. To extract proteins, the pellet was resuspended in lysis buffer and sonicated for 20 cycles on ice. Afterward, the lysate was gathered by centrifuging at 17,467 x g for 30 minutes at 4 °C.

Subsequently, the lysate was mixed with 20% trichloroacetic acid in acetone, vortexed, and chilled on ice. Post-centrifugation, the supernatant was removed, and the pellet was rinsed with acetone, centrifuged, and washed once more with 80% acetone in water. The protein pellet was air-dried and dissolved in 50 mM Tris buffer. For protein purification, SpeedBead Carboxylate-Modified E3 and E7 (Sera-Mag, USA) were used, and the EZQ protein assay kit (Invitrogen, USA) was employed for protein concentration quantification. The sample then underwent reduction, alkylation, and trypsin digestion.

After the digestion process, the sample was diluted, passed through a molecular sieve, and exposed to solid-phase extraction. Proteins were eluted and concentrated using a speed vacuum, and a 10 µL aliquot from each sample was analyzed via nano LC-MS/MS. A NanoLC 400 UPLC system (Eksigent, USA) was utilized for sample desalting and separation, while a TripleTOF 6600 Quadrupole-Time-of-Flight mass spectrometer (Sciex, USA) performed mass spectrometry analysis.

The resulting data were searched against a database established by metagenomics results of the samples using MetaProteomeAnalyzer version 3.4^14^. The X-tandem was selected as the primary peptides and protein identification search engine^15^. The parameters were as follows: precursor and fragments tolerance set as 15 ppm; the cysteine alkylation set as iodoacetamide; the trypsin as the digestion enzyme with up to one mis-cleavage; a 1% false discovery rate was set as the filter of the final global protein groups. The resulting group file exported from X-tandem was converted to MetaProteomeAnalyzer for metaproteomics analysis and clustering. Information on enzymes regulated by regulons comes from RegulonDB v12.0^16^.

**S2. RESULTS**

**S2.1. Treatment performances under different aeration conditions**

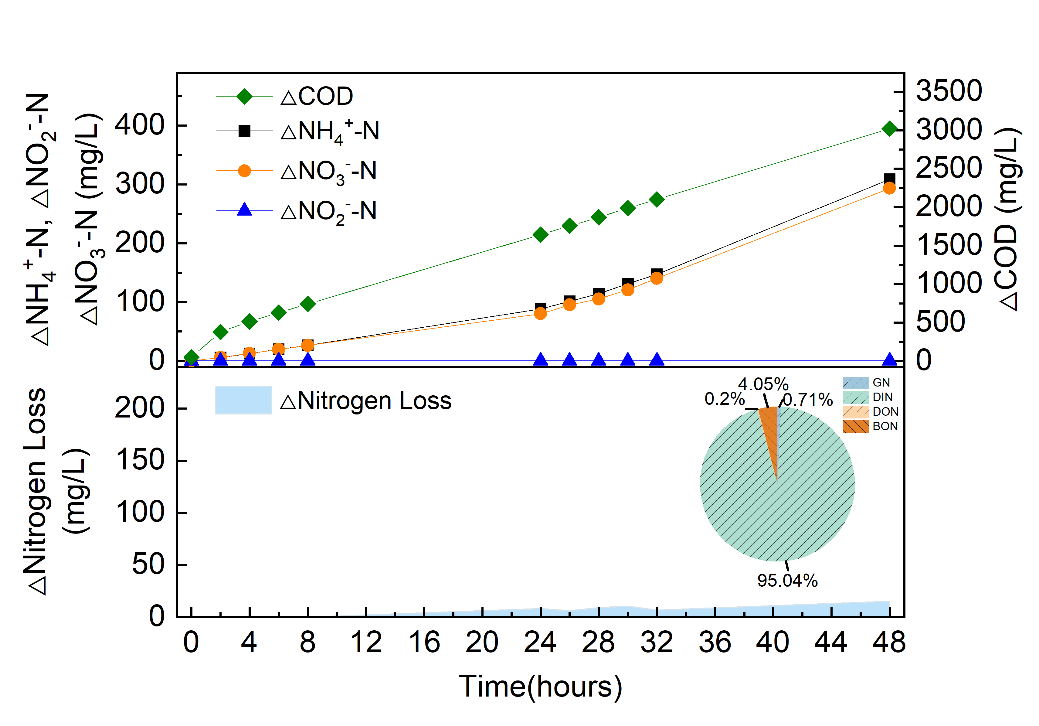

**Figure S1.** The concentration of carbon and inorganic nitrogen in the bioreactor under CA2 condition

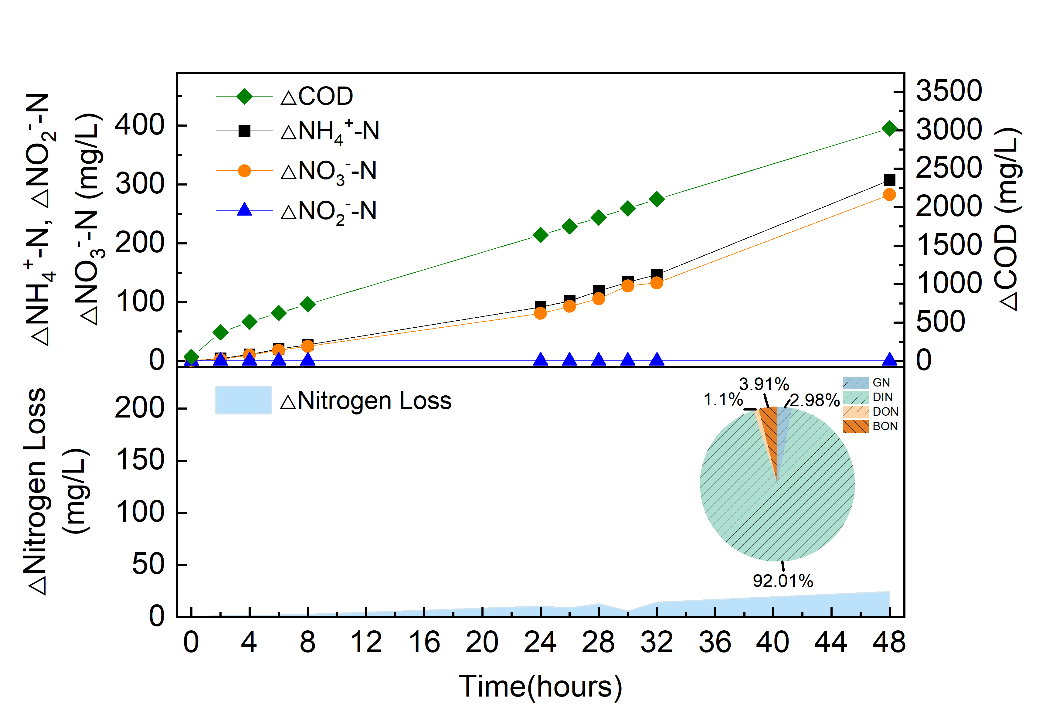

**Figure S2.** The concentration of carbon and inorganic nitrogen in the bioreactor under CA8 condition

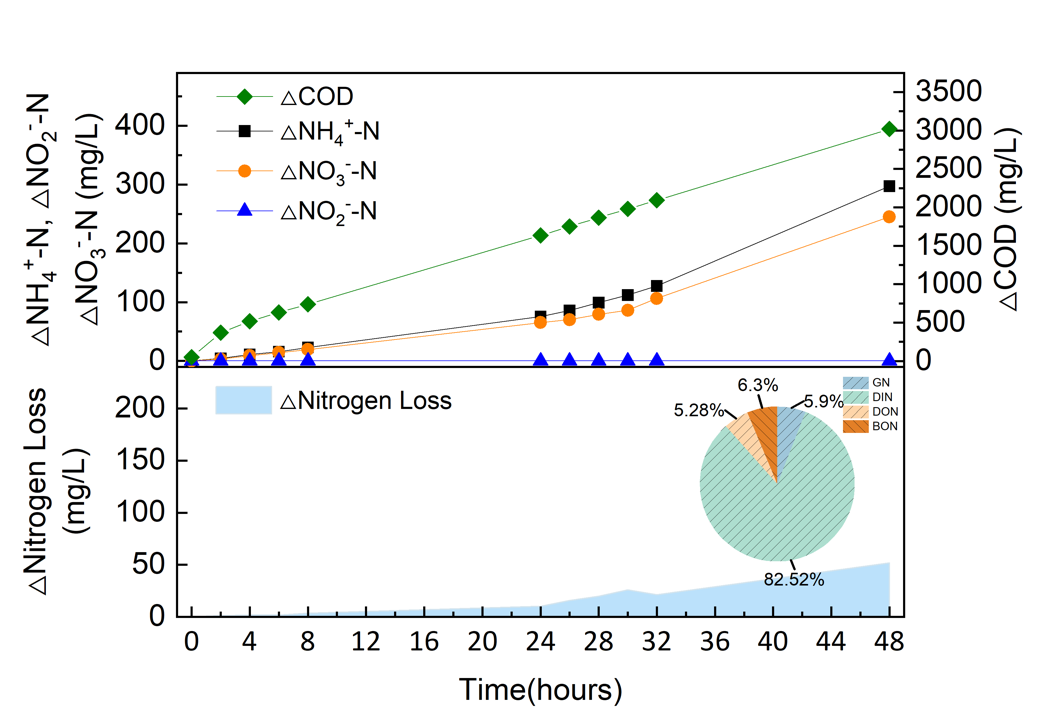

**Figure S3.** The concentration of carbon and inorganic nitrogen in the bioreactor under CP2 condition

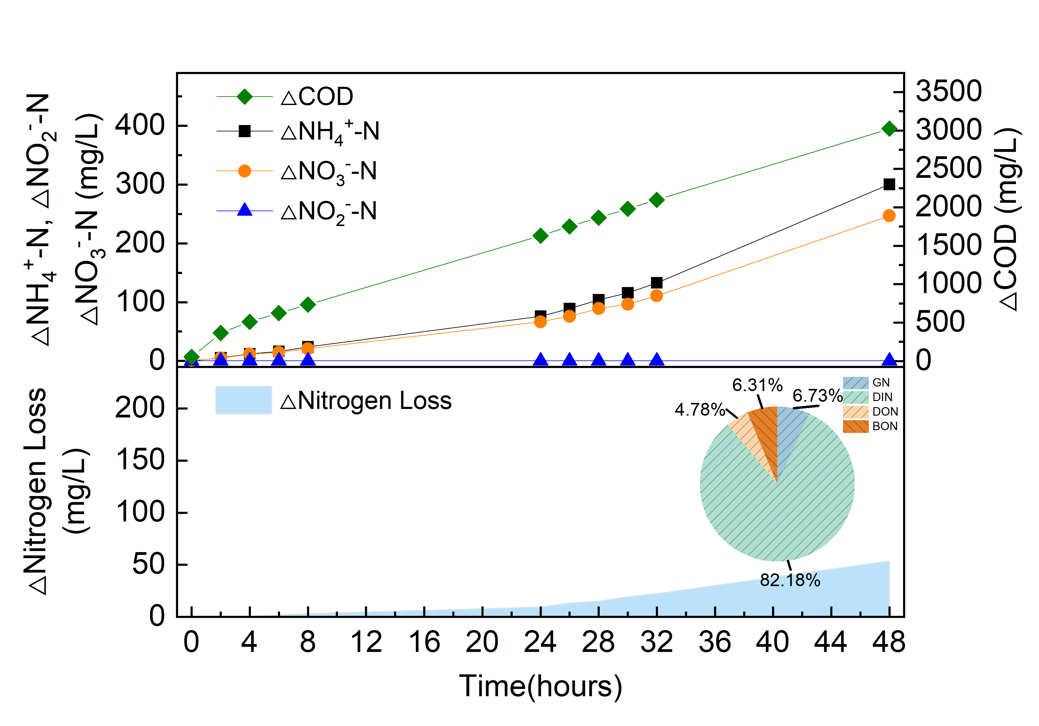

**Figure S4.** The concentration of carbon and inorganic nitrogen in the bioreactor under CP8 condition

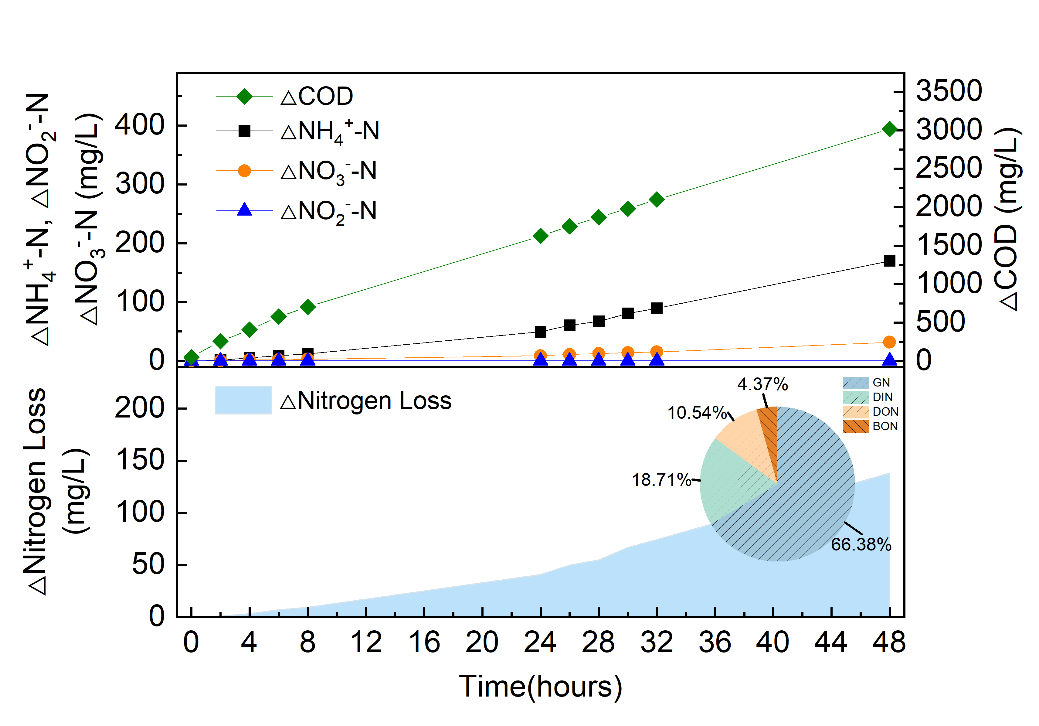

**Figure S5.** The concentration of carbon and inorganic nitrogen in the bioreactor under IP2 condition

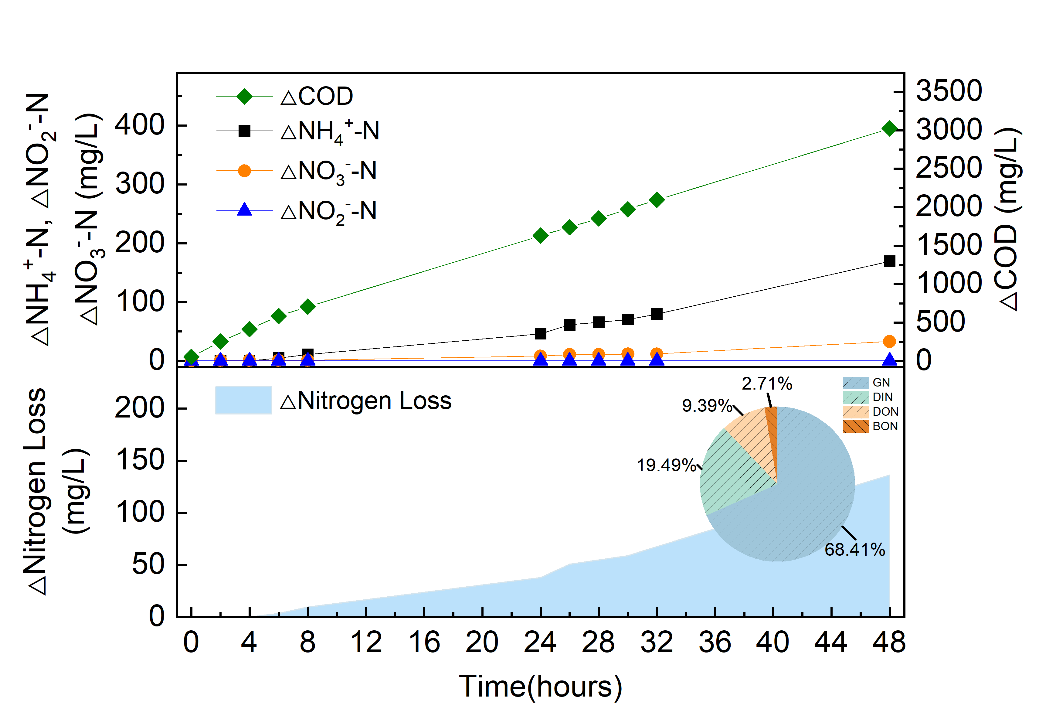

**Figure S6.** The concentration of carbon and inorganic nitrogen in the bioreactor under IP8 condition

**S2.2. Dissolved N_2_O accumulation dynamics during 48 hours of exposure to different aeration conditions**

**Table S3.** Phase-specific dissolved N_2_O accumulation rates and confidence intervals for various aeration conditions

| **Condition** | **Phase** | **Time Range (h)** | **N_2_O Accumulation Rate (mg/L/h)** | **Standard Error (mg/L/h)** | **95% Confidence Interval (mg/L/h)** |
| --- | --- | --- | --- | --- | --- |
| CA2 | Phase I | 0-9.15124 | 0.0386 | 4.97486×10^−5^ | [0.03850, 0.03871] |
| CA2 | Phase II | 9.15124-40.29318 | -0.00689 | 1.11039×10^−5^ | [-0.00691, -0.00687] |
| CA2 | Phase III | 40.29318-48 | -0.00295 | 1.03367×10^−4^ | [-0.00315, -0.00275] |
| CA8 |  | 0-48 | -0.00159 | 2.97062×10^−6^ | [-0.00160, -0.00158] |
| CP2 | Phase I | 0-9.43405 | 0.06357 | 1.05554×10^−4^ | [0.06336, 0.06378] |
| CP2 | Phase II | 9.43405-31.24229 | -0.02339 | 3.00661×10^−5^ | [-0.02345, -0.02333] |
| CP2 | Phase III | 31.24229-48 | -0.00298 | 4.56785×10^−5^ | [-0.00307, -0.00289] |
| CP8 | Phase I | 0-7.75595 | 0.06767 | 2.69046×10^−4^ | [0.06716, 0.06818] |
| CP8 | Phase II | 7.75595-20.83103 | -0.03928 | 1.23107×10^−4^ | [-0.03952, -0.03904] |
| CP8 | Phase III | 20.83103-48 | -2.53266×10^−5^ | 4.16114×10^−5^ | [-0.00010, 0.00005] |
| IP2 | Phase I | 0-16.93672 | 0.03394 | 6.28413×10^−5^ | [0.03382, 0.03406] |
| IP2 | Phase II | 16.93672-36.28074 | -0.02224 | 5.16004×10^−5^ | [-0.02234, -0.02214] |
| IP2 | Phase III | 36.28074-48 | -0.00753 | 1.12148×10^−4^ | [-0.00775, -0.00731] |
| IP8 | Phase I | 0-11.04823 | 0.00765 | 7.50778×10^−5^ | [0.00748, 0.00783] |
| IP8 | Phase II | 11.04823-34.20828 | -0.00529 | 2.47674×10^−5^ | [-0.00533, -0.00525] |
| IP8 | Phase III | 34.20828-48 | -0.00209 | 5.50477×10^−5^ | [-0.00220, -0.00198] |

**S2.3. N_2_O consumption rates after 48 hours of exposure to different aeration conditions**

**Table S4.** In-situ N_2_OR activity of microbial systems under different aeration conditions after 48 hours

| **Condition** | **In-situ N_2_OR Activity (mgN_2_O-N/L/min)** | **Standard Error (mgN_2_O-N/L/min)** | **R-Square** |
| --- | --- | --- | --- |
| CA2 | 0.3330 | 4.431×10^−4^ | 0.99832 |
| CA8 | 0.0828 | 4.200×10^−4^ | 0.98496 |
| CP2 | 0.3120 | 1.906×10^−3^ | 0.97535 |
| CP8 | 0.6396 | 1.162×10^−3^ | 0.99418 |
| IP2 | 1.2306 | 4.733×10^−3^ | 0.99135 |
| IP8 | 4.1874 | 9.990×10^−3^ | 0.99634 |

**S2.4. The enzyme expression related to nitrogen transformation in the microbial system before and after 48 hours of exposure to different aeration conditions**

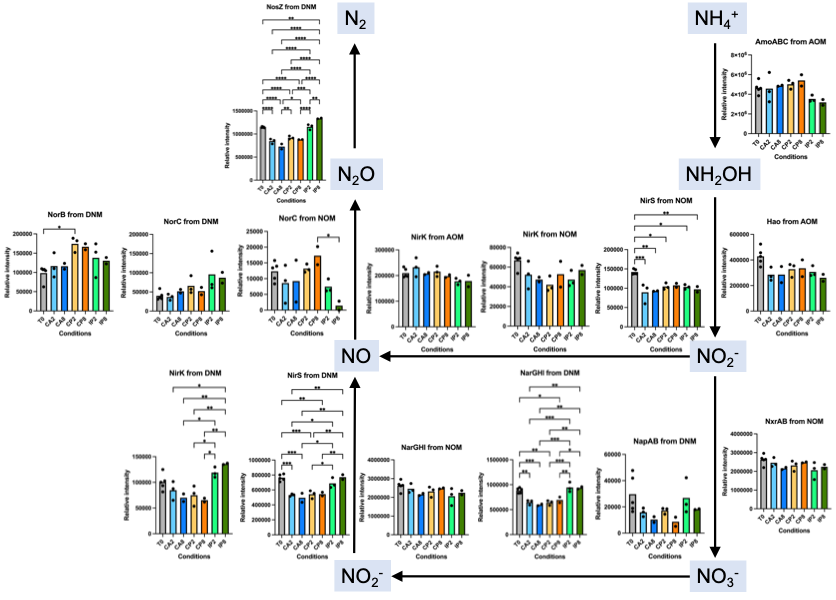

**Figure S7.** Abundance of key nitrogen cycle enzymes from ammonia-oxidizing microorganisms (AOM), nitrite-oxidizing microorganisms (NOM), and denitrifying microorganisms (DNM) in microbial systems at the start of the reaction (T0) and under six different aeration conditions

**S2.5. The nitric oxide concentrations under different aeration conditions**

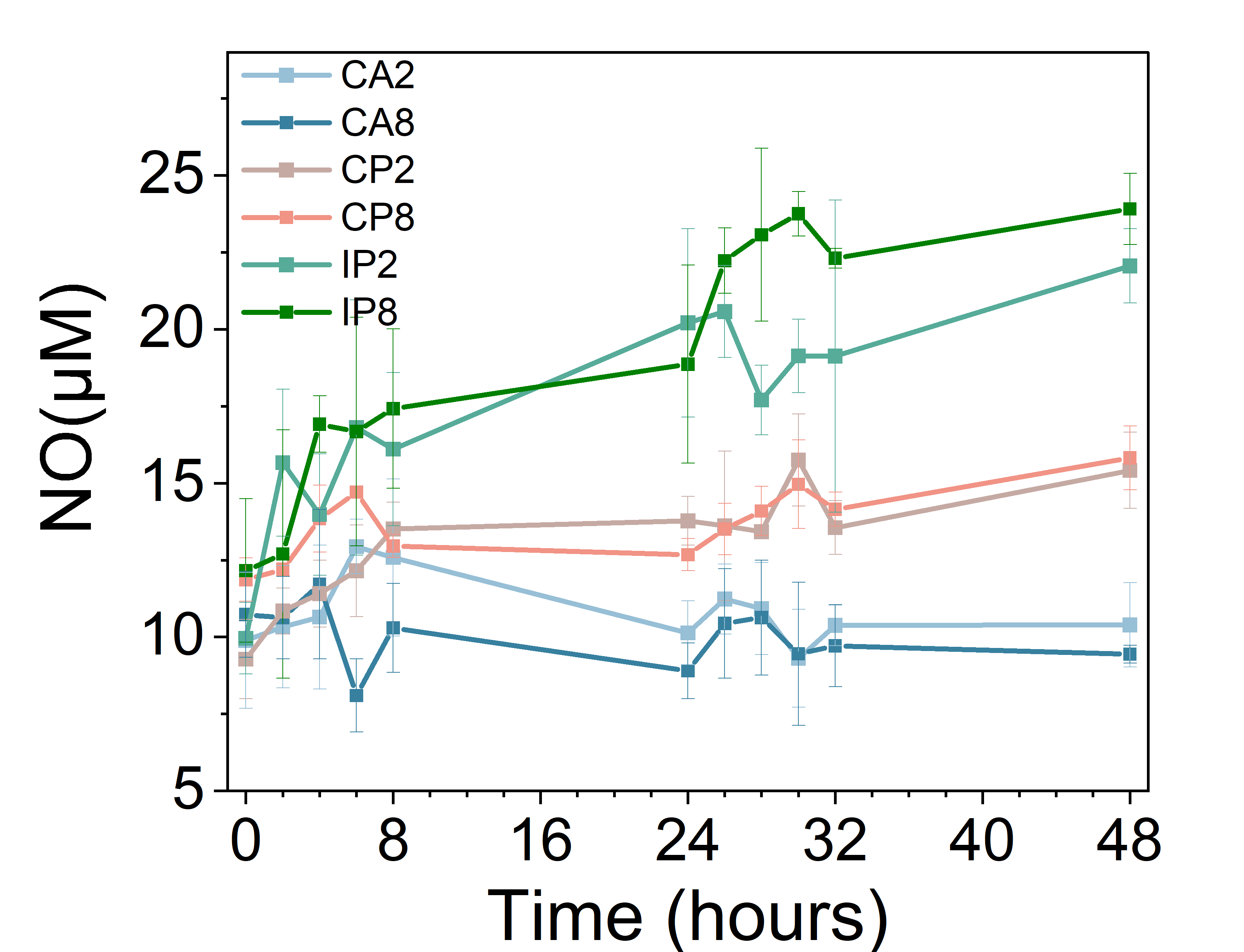

### Figure S8. The concentration of NO under different aeration conditions: constant aeration (CA), continuous perturbation (CP), and intermittent perturbation (IP). The error envelope represents standard deviations (technical duplication results for biological triplicates; n=6).

**S2.6. The enzyme expression from *Methylobacillus* after 48 hours of exposure to different aeration conditions**

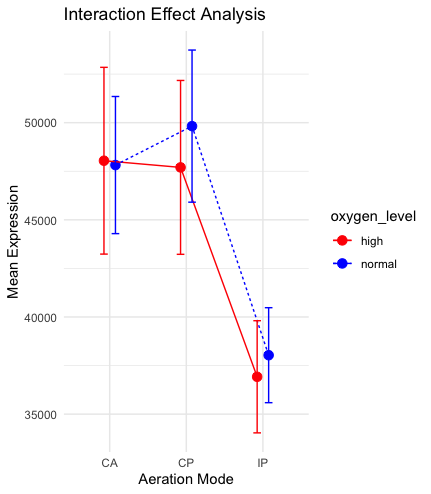

**Figure S9.** The mean expression levels and standard errors for enzyme expression from *Methylobacillus* under different aeration conditions by two-way ANOVA analysis (normal oxygen level conditions: CA2, CP2, IP2; high oxygen level conditions: CA8, CP8, IP8)

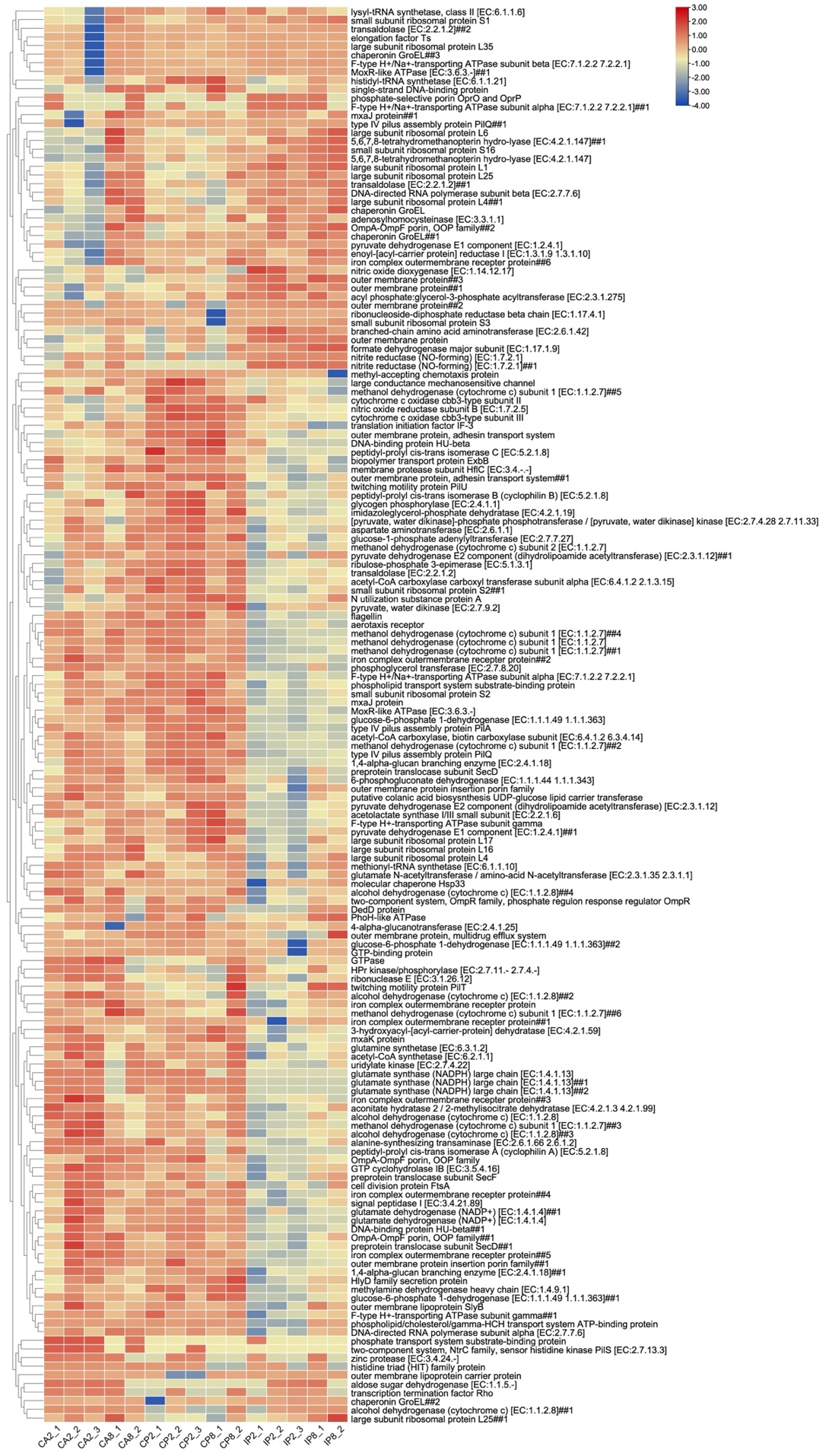

**Figure S10.** The enzymes in *Methylobacillus* that exhibited significant differences in abundance under different aeration modes

**S2.7. The analysis for enzyme expression from *Hyphomicrobium* after 48 hours of exposure to different aeration conditions**

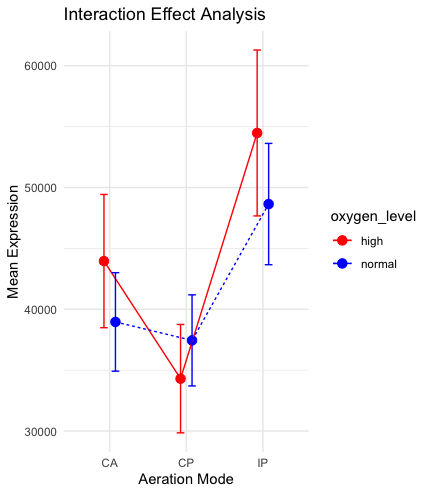

**Figure S11.** The mean expression levels and standard errors for enzyme expression from *Hyphomicrobium* under different aeration conditions by two-way ANOVA analysis (normal oxygen level conditions: CA2, CP2, IP2; high oxygen level conditions: CA8, CP8, IP8)

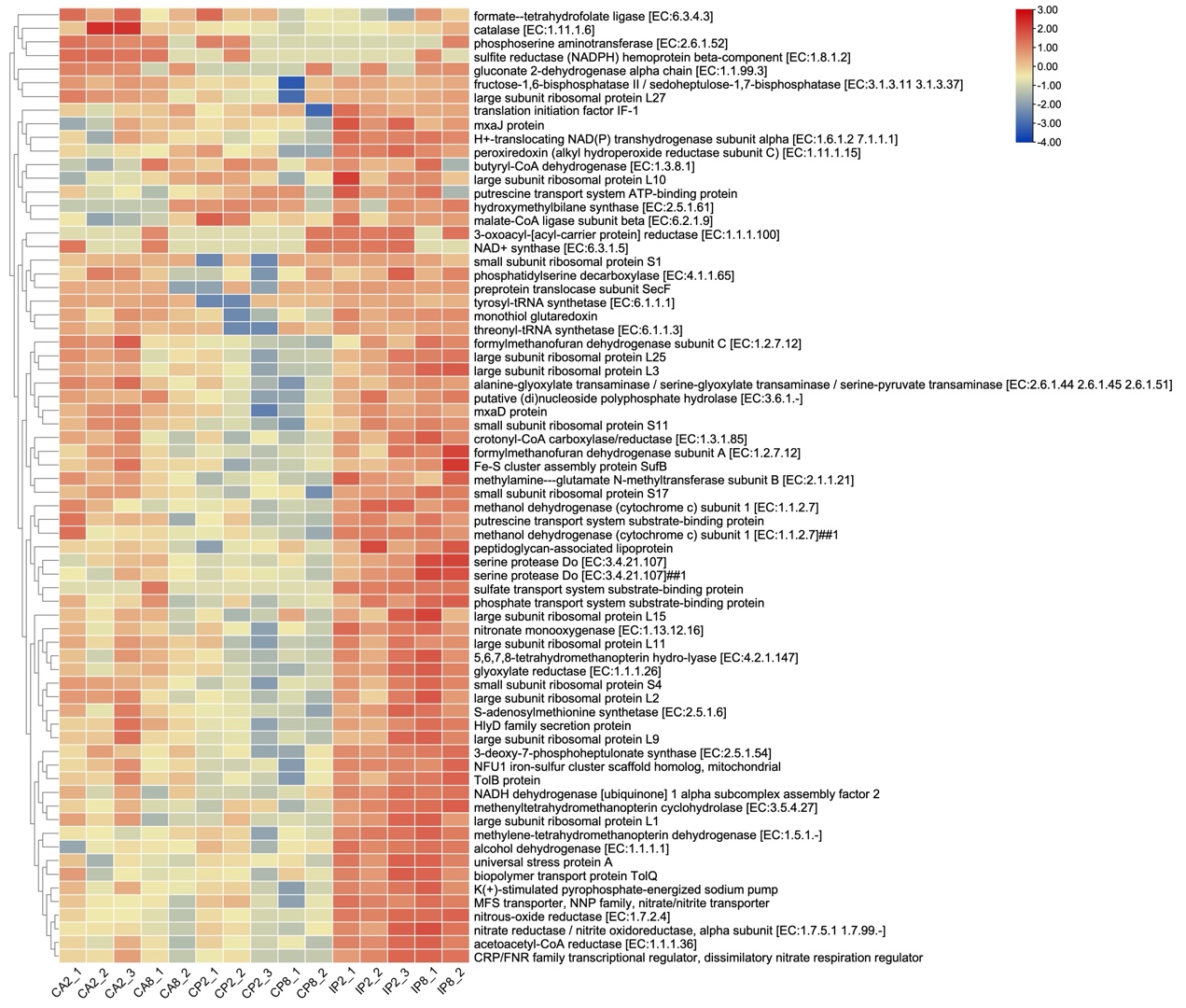

**Figure S12.** The enzymes in *Hyphomicrobium* that exhibited significant differences in abundance under different aeration modes
